# Antiviral Effect of Hyunggaeyungyo-tang on A549 Cells Infected with Human Coronavirus

**DOI:** 10.1101/2021.06.02.446680

**Authors:** Seo-Young Won, In-Chan Seol, Ho-Ryong Yoo, Yoon-Sik Kim

## Abstract

**Background:** Herbal medicine is widely recommended to treat viral infectious diseases. Over 123,000,000 individuals have been infected with the coronavirus since a worldwide pandemic was declared in March 2020. We conducted this research to confirm the potential of herbal medicine as a treatment for coronavirus.

**Methods:** We infected the A549 cell line with beta coronavirus OC43 then treated with 100 μg/mL Hyunggaeyungyo-tang (HGYGT) or distilled water with a control of HGYGT. We measured the mRNA expression levels of pro-inflammatory cytokines and interferon stimulated genes (ISGs) to confirm the effectiveness of HGYGT upon coronavirus infection.

**Results:** We found the effects of HYGYT decrease the expression level of pPKR, peIF2α, IFI6, IFI44, IFI44L, IFI27, IRF7, OASL and ISG15 when administered to cells with coronavirus infection. The expressions of IL-1, TNF-α, COX-2, NF-κB, iNOS and IKK mRNA were also significantly decreased in the HGYGT group than in the control group.

**Conclusion:** Through the reduction of the amount of coronavirus RNA, our research indicates that HGYGT has antiviral effects. The reduction of IKK and iNOS mRNA levels indicate that HGYGT reduces coronavirus RNA expression and may inhibits the replication of coronavirus by acting on NF-kB/Rel pathways to protect oxidative injury. In addition, decreases in mRNA expression levels of pro-inflammatory cytokines indicate that the HGYGT may relieve the symptoms of coronavirus infections.

## 1. Introduction

The number of infectious diseases caused by coronaviruses has recently increased. The severe acute respiratory syndrome coronavirus 1 (SARS-CoV-1) led to a pandemic in 2002, the Middle East respiratory syndrome (MERS)-CoV in 2012, and the SARS-CoV-2 in 2019. Coronaviruses are divided into four types: alpha, beta, gamma, and delta. Human CoV (HCoV) belongs to the genera *Alphacoronavirus* and *Betacoronavirus*. The 229E and NL63 types are Alphacoronaviruses. OC43, HKU1, SAR-CoV-1, MER-CoV, and SAR-CoV-2 are Betacoronaviruses. A novel CoV (2019-nCoV; SAR-CoV-2), identified in January 2020, results in respiratory symptoms and has spread worldwide. The recent coronavirus infection, named coronavirus disease 2019 (COVID-19) by the World Health Organization, has led to a global pandemic.

The COVID-19 Korean Medicine Treatment Recommendation, written based on the National Health Commission of the People’s Republic of China’s treatment plan, recommends herbal medicine treatment at the appropriate time. Some guidelines recommend that uninsured herbal extracts are administered for symptomatic relief. However, in Korea, uninsured herbal extracts are generally perceived to be ineffective compared to packed herbal medicines. Therefore, we determined the effectiveness of an uninsured herbal extract for the treatment of COVID-19.

Hyunggaeyungyo-tang (HGYGT) is an herbal medicine composed of 13 herbs (Table 1). HGYGT is widely prescribed in patients with anhidrosis, excessive thirst, excessive sputum, fever, or sore throat. It has been reported that 88% of patients with COVID-19 had fever, 65% had dyspnea, and 60% had a cough [1]. HGYGT is prescribed for these symptoms. We confirmed the antiviral effect of HGYGT using a cell line infected with OC43, a relatively safe Betacoronavirus.

**Table 1.** Composition of HGYGT.

| Scientific name | Amount (g) |
| --- | --- |
| *Schizonepeta tenuifolia* Briquet (KP) | 0.5 |
| *Scutellaria baicalensis* Georgi (KP) | 0.5 |
| *Saposhnikovia divaricata* Schischkin (KP) | 0.5 |
| *Angelica gigas* Nakai (KP) | 0.5 |
| *Poncirus trifoliata* Rafinesque (KP) | 0.5 |
| *Paeonia lactiflora* Pallas (KP) | 0.5 |
| *Glycyrrhiza uralensis* Fischer (KP) | 0.5 |
| *Cnidium officinale* Makino (KP) | 0.5 |
| *Angelica dahurica* Bentham et Hooker f (KP) | 0.83 |
| *Gardenia jasminoides* Ellis (KP) | 0.5 |
| *Platycodon grandiflorum* A. De Candolle (KP) | 0.83 |
| *Forsythia suspensa* Vahl (KP) | 0.5 |
| *Bupleurum falcatum* Linné (KP) | 0.83 |

HGYGT: Hyunggaeyungyo-tang; KP: Korean Pharmacopeia

Protein kinase RNA-activated (PKR) is one of four mammalian serine-threonine kinases that primarily react to double-stranded RNA (dsRNA) during a viral infection and phosphorylate the eukaryotic initiation factor 2α (eIF2α) translation initiator. The phosphorylation of eIF2α inhibits the translation of viral mRNAs, ceasing viral replication [2]. In addition, dsRNA is a potent inducer of type I IFN, which produces interferon stimulated genes (ISGs) and forms a protective antiviral state [3]. Virus infection produces reactive oxygen specifications (ROS), which causes virus replication and oxidative injury. [4] We identified the presence of immune-related proteins such as PKR, phosphorylated PKR(pPKR), eIF2 and phosphorylated eIF2α(peIF2α) using immunocytochemistry and measured the expression of proinflammatory cytokines and ISGs to confirm and understand the antiviral effects of HGYGT in A549 cells infected with human coronaviruses.

## 2. Materials and Methods

### 2.1. Reagents and Instruments

The HGYGT used in this experiment was purchased from Hanpoong Pharm.(Jeonju, Korea). The amount of HGYGT’s validity was calculated to control the concentration of the active ingredient. HGYGT was diluted with physiological saline to 100 μg/mL.

The reagents used are TRIsure (Bioline, England), SYBR (Bioline, England), Reverse transcriptase (Thermo Fisher Scientific, USA), dNTP (Takara, Japan), DNase (Takara, Japan), RNase inhibitor (Takara, Japan), protease inhibitor (Takara, Japan), primary antibody (Cell signaling technology, USA), secondary antibody (Thermo Fisher Scientific, USA), Sodium acetate (Invitrogen, USA), Ethanol (Sigma, Germany).

The devices used are RT-PCR (Thermo Fisher Scientific, USA), micro reader (Thermo Fisher Scientific, USA), PCR machine (Thermo Fisher Scientific, USA), chemidoc (Thermo Fisher Scientific, USA).

### 2.2. Cell culture

A549 cells were cultured in RPMI medium (Welgene, Korea) and 10% fetal bovine growth serum (RMbio, USA). The cells were cultured at 37°C in a humidified atmosphere containing 5% CO_2_.

### 2.3. Virus infection

OC43 (1.0 MOI) was added to 5 × 10^5^ A549 cells and the cells were incubated for three days. Cells were then treated with HGYGT (100 μg/mL) or deionized, distilled water (DW). The cells were cultured for 72 h prior to RNA extraction.

### 2.4. SRB Viability Assay

Two days after drug treatment, the cells were fixed during 1hr with 10% TCA solution, washed twice with DPBS, and dried at room temperature. The cells were stained with a 0.05% SRB solution. The stained cells were washed 4 times with 1% acetic acid and dried at room temperature. After washing four times with DPBS, the cells were then reconstituted 1hr at room temperature using 10 mM Tris (pH = 10.5). The supernatant of Tris washed samples are analyzed with a microplate reader at an absorbance of 510 nm.

### 2.5. RNA Extraction

The total RNA was extracted using TRIsure (Bioline, England). Prepared cells were distributed into an 8 pi medium consisting of 1 × 10^6^ cells on 100 pi plates. A549-OC43 cells (OC43 group) and A549-OC43 cells treated with HGYGT (HGYGT group) were cultured for 72 h. All prepared groups were incubated for 5 min at room temperature before treatment with 1 mL of TRIsure per 5 × 10^5^ cells. The lysate was passed several times with a pipette tip. After incubation for 5 min at room temperature, chloroform was added, and the mixture was vortexed for 15 min then centrifuged for 1 h at 15,000 rpm. After the supernatant was discarded, the pellet was blended with cold isopropyl alcohol to precipitate the RNA. The sample was incubated at room temperature for 10 min and centrifuged at 12,000 rpm at 4°C for 10 min. The pellet was washed using 75% ethanol, air-dried, and dissolved in PCR water. To remove the genetic DNA, purified nucleic acids were treated with DNA enzyme I (Takara, Japan). RNA was reverse transcribed using RevertAid reverse transcriptase (Thermo Fisher Scientific, USA). The sample was centrifuged for 10 min at 12,000 rpm and room temperature and the supernatant was discarded. After it was washed twice with 75% ethanol and dried, the pellet was dissolved in DW.

### 2.6. cDNA Synthesis and RT-PCR

For reverse transcription (RT) reactions, PCR was performed with 800 ng of total RNA prepared with 1 μL of random primer. The sample was denatured at 65°C for 5 min, then the temperature dropped to 4°C before 4 μL of a 10 mM dNTP mixture and 1 μL of reverse transcriptase were added with 1 μL RNase inhibitor (20 U/μL) and 4 μL 5×RT buffer (250 mM Tris-HCl, pH = 8.3, 375 mM KCl, 15 mM MgCl_2_). PCR was performed at 25°C for 10 min, at 42.5°C for 60 min, and at 70°C for 10 min. RT-PCR (Bio-Rad, USA) was performed using synthesized cDNA, forward/reverse primers, and the SensiFAST SYBR Lo-Rox Kit (Bioline, England). The sequences of primers used in the experiment are presented in Table 2. The forward/reverse primer mixture (2 μL at 3 μM) was combined with 7.5 μL SYBR, 4.5 μL DW, and 1 μL cDNA. In addition, pre-denaturation was performed for 40 cycles at 5 min each at 95°C, 40 cycles at 95°C, and 1 min at 60°C. The Cq values were calculated.

**Table 2.**
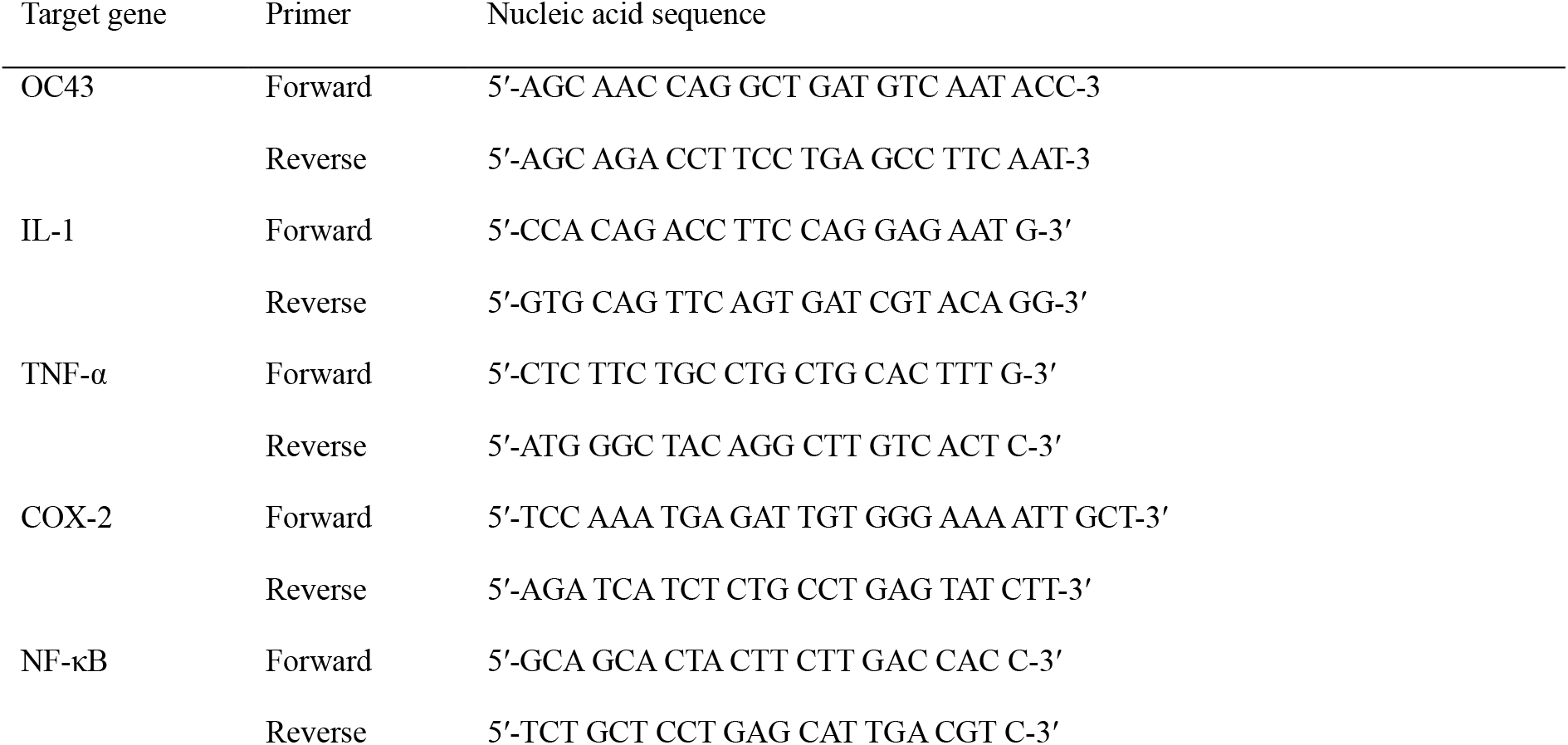
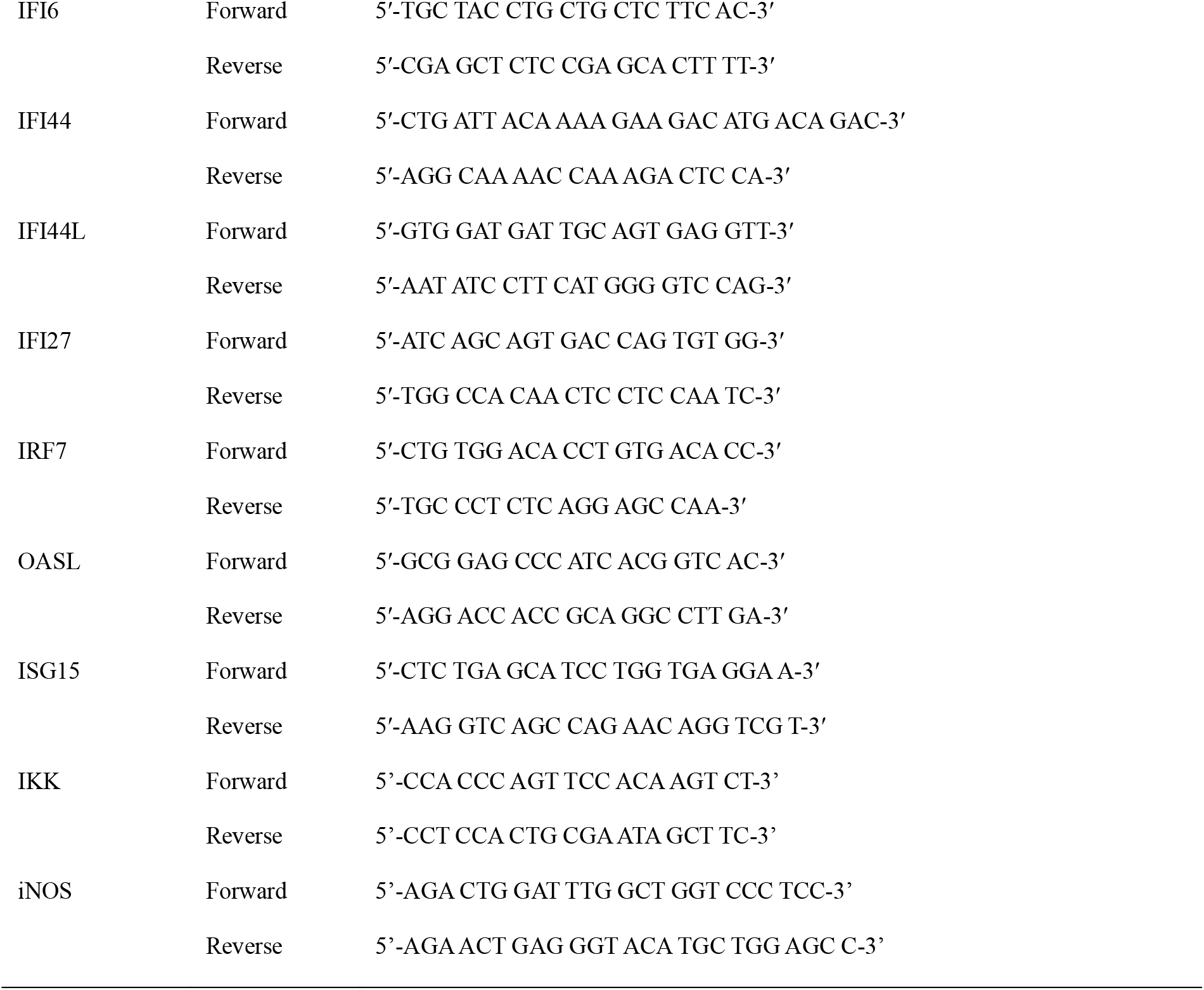
Primer Sequences used in RT-PCR.

### 2.7. Immunocytochemistry

The cells were washed twice with PBS and fixed in 4% paraformaldehyde for 15 min at room temperature. Then, the cells were permeabilized with 0.1% Triton X-100 in BSA buffer and blocked for 1 h at room temperature. Cells were incubated with primary antibodies diluted with 1% BSA for two h then washed four times with 0.1% (v/v) Tween-20 in PBS and incubated with Alexa Fluor-conjugated secondary antibodies. The cells were imaged using a Zeiss LSM 760 confocal microscope with a C-Apochromat 20x objective lens (NA = 1.40). PKR and pPKR primary antibodies were purchased from Santa Cruz Biotechnology; eIF2α and peIF2α antibodies were obtained from Cell Signaling Technology.

### 2.8. Statistical Analysis

Variables are expressed as mean ± SEM. Unpaired one-tailed Student’s t-tests were used to compare the data. Statistical significance was set at p < 0.05.

## 3. Results

### 3.1. OC43 mRNA expression

The OC43 mRNA expression was significantly higher in the OC43 group (1.019 ± 0.1413) than in the HGYGT group (0.4205 ± 0.07611) (p < 0.05) (Table 3, Figure 1).

**Table 3.** mRNA Expression.

| mRNA | Expression in control group | Expression in OC43 group.<br>(p-value vs. control) | Expression in HGYGT group<br>(p-value vs. OC43) |
| --- | --- | --- | --- |
| OC43 | | $1.019 \pm 0.1413$ | $0.4205 \pm 0.07611$<br>*0.0102 |
| IL-1 | $1.002 \pm 0.04936$ | $27.75 \pm 1.868$<br>****<0.0001 | $21.92 \pm 0.6333$<br>*0.0209 |
| TNF-α | $1.054 \pm 0.2352$ | $225.8 \pm 24.08$<br>***0.0004 | $166.7 \pm 8.045$<br>*0.0401 |
| COX-2 | $1.021 \pm 0.1516$ | $10.89 \pm 0.8811$<br>***0.0002 | $8.609 \pm 0.3086$<br>*0.0356 |
| NF-κB | $1.009 \pm 0.09722$ | $6.482 \pm 1.039$<br>**0.0032 | $3.807 \pm 0.07861$<br>*0.0311 |
| IFI6 | $1.026 \pm 0.1712$ | $150.5 \pm 15.18$<br>***0.0003 | $84.61 \pm 7.541$<br>**0.0089 |
| IFI44 | $1.071 \pm 0.2847$ | $1413 \pm 169.2$<br>***0.0006 | $761.7 \pm 49.3$<br>*0.0105 |
| IFI44L | $1.009 \pm 0.09376$ | $137.1 \pm 4.453$<br>****<0.0001 | $98.37 \pm 2.426$<br>***0.0008 |
| IFI27 | $1.002 \pm 0.03935$ | $506.2 \pm 48.8$<br>***0.0002 | $328 \pm 29.63$<br>*0.0177 |
| IRF7 | $1.009 \pm 0.09585$ | $7.599 \pm 0.6307$<br>***0.0002 | $5.222 \pm 0.7767$<br>*0.0382 |
| OASL | $1.003 \pm 0.05799$ | $86.96 \pm 1.575$<br>****<0.0001 | $59.74 \pm 2.015$<br>***0.0002 |
| ISG15 | $1.004 \pm 0.06054$ | $71.27 \pm 4.874$<br>****<0.0001 | $48.62 \pm 4.098$<br>*0.0118 |
| IKK | $1.004 \pm 0.06449$ | $2.6 \pm 0.0503$<br>****<0.0001 | $1.566 \pm 0.1109$<br>****<0.0001 |
| INOS | $1.002 \pm 0.04371$ | $2.195 \pm 0.05391$<br>****<0.0001 | $1.372 \pm 0.02985$<br>****<0.0001 |

Values are presented as mean ± SEM (n=3), (\* ; $p < 0.05$ , \*\* ; $p < 0.01$ , \*\*\* ; $p < 0.001$ , \*\*\*\* ; $p < 0.0001$ )

**Fig 1.**
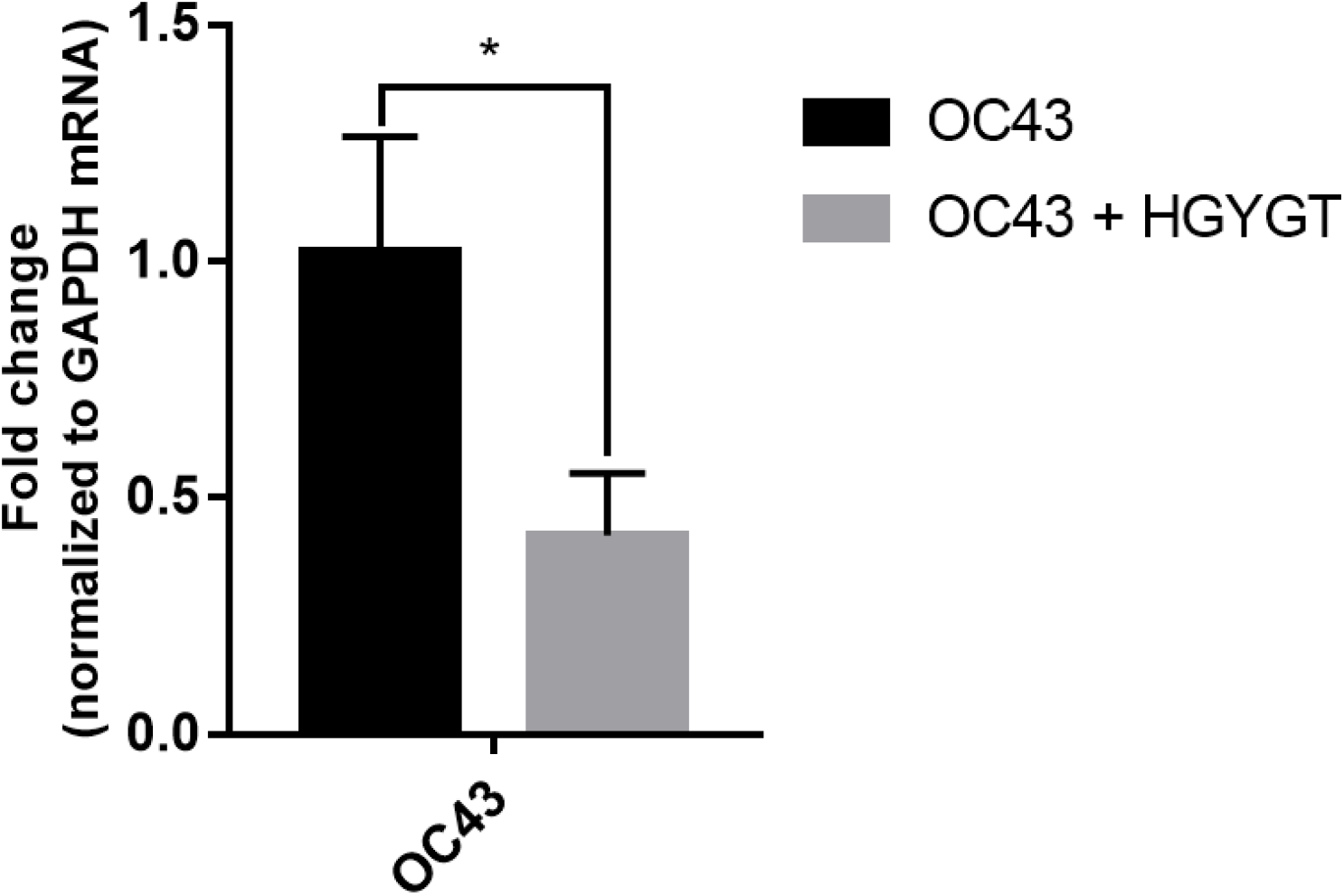
OC43 mRNA expression in A549 cells. mRNA expression is measured using quantitative real-time PCR and presented as mean ± S.D. (n= 3). OC43: cells treated with OC43; OC43 + HGYGT: cells treated with OC43 and HGYGT (100 μg/mL). (*; p<0.05)

### 3.2. Cell Visualizing

Though the difference was not significant, the expression of PKR was higher in the cells infected OC43 than cells treated with both OC43 and HGYGT. The expression of pPKR was significantly increased in the cells treated only with OC43 compared to the expression in cells treated with both OC43 and HGYGT. The expression of pPKR in the HGYGT group and uninfected A549 cells was not significantly different (Figure 2). The expression of peIF2α was significantly higher in the control group than in the cells that were infected with OC43 and treated with HGYGT (Figures 3 and 4).

**Fig 2.**
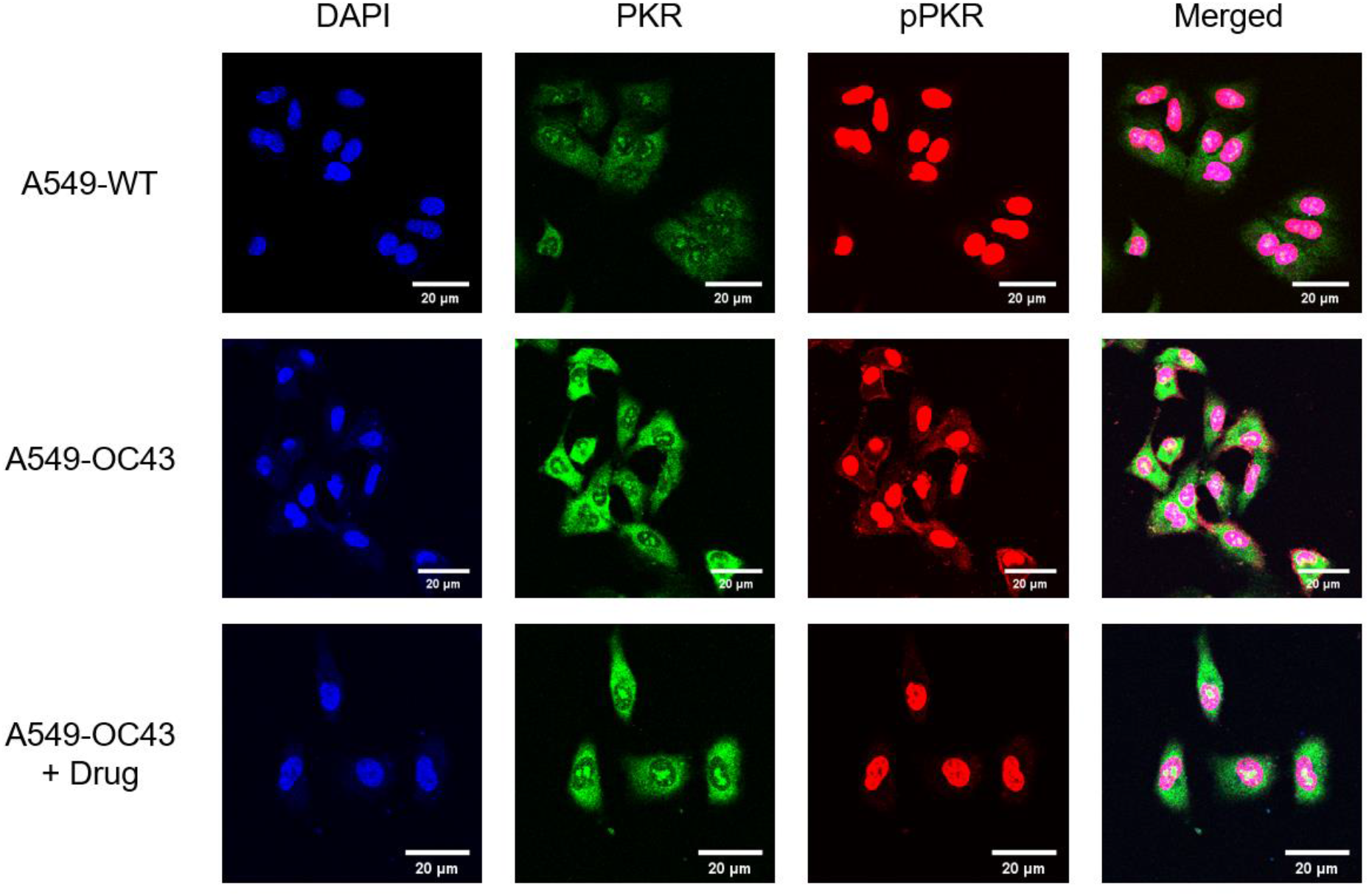
Visualization of PKR, pPKR expression.

**Fig 3.**
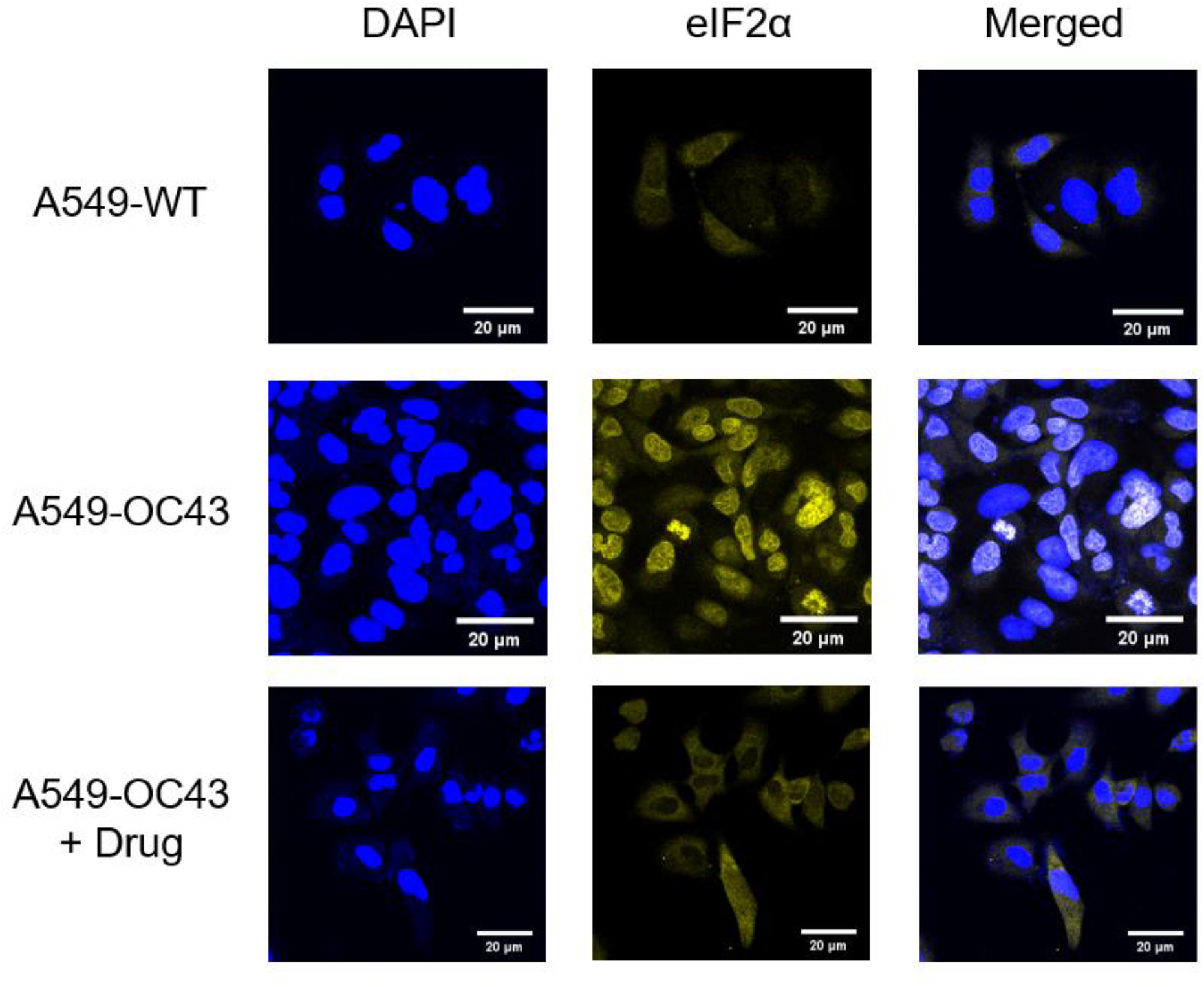
Visualization of eIF2α expression.

**Fig 4.**
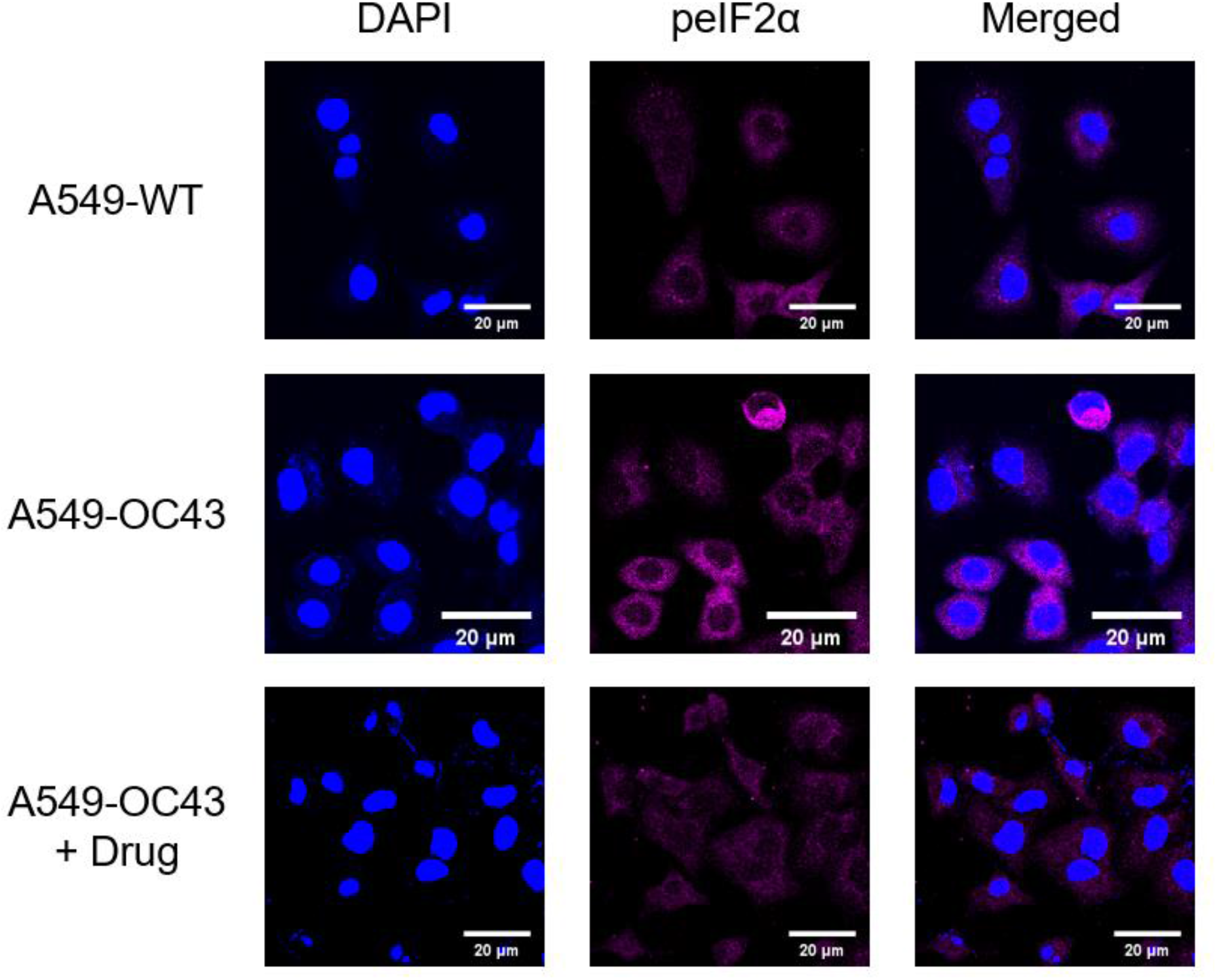
Visualization of peIF2α expression.

### 3.3. Cell viability

The viabilities of cells treated with 100 μg/mL, 200 μg/mL, 300 μg/mL, 400 μg/mL and 500 μg/mL HGYGT were 100.891%, 97.4155%, 98.3408%, 100.627%, 103.914%, respectively. There are no side effects in drug treatment (Figure 5).

**Fig 5.**
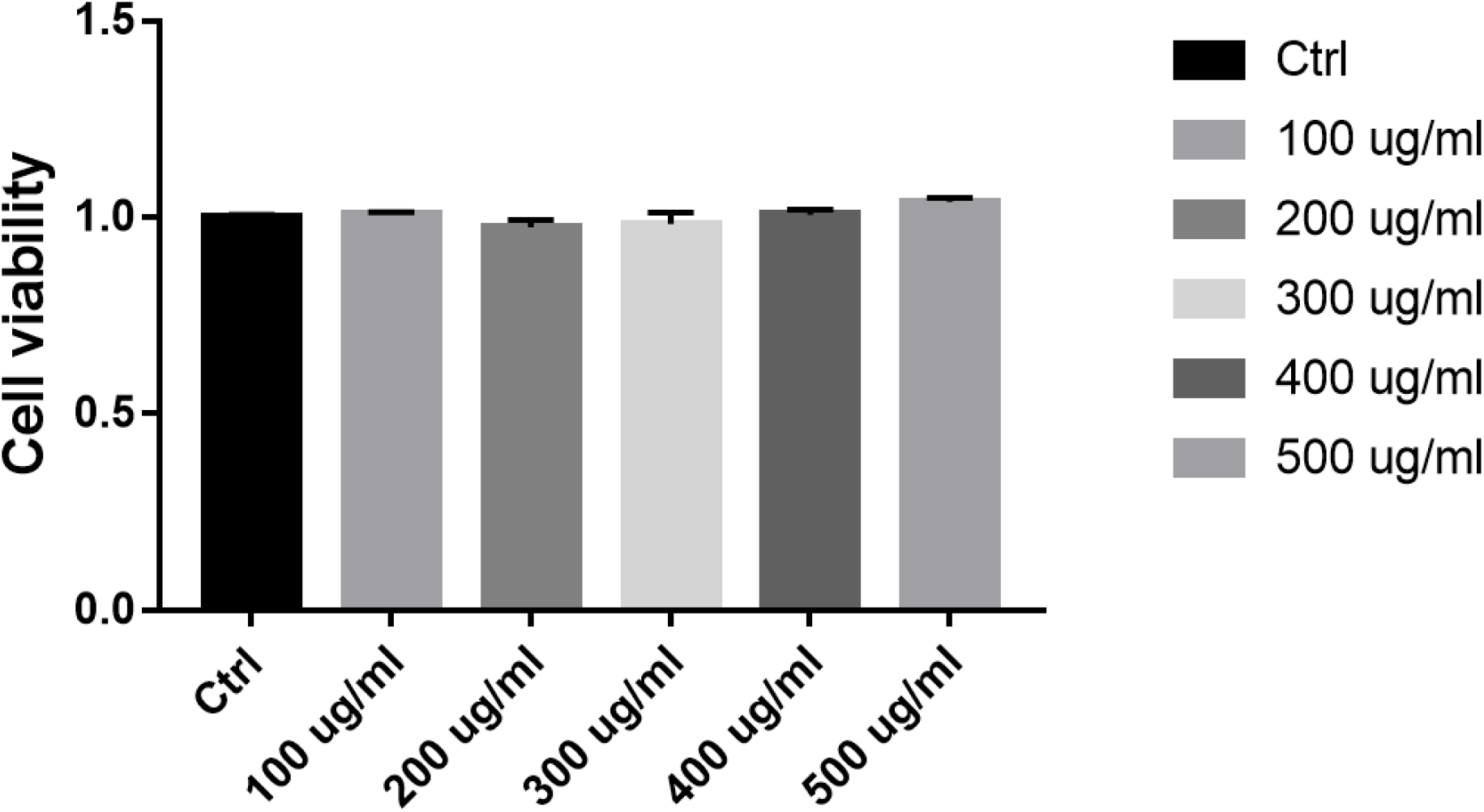
SRB assay of HGYGT in A549 cells.

### 3.4. mRNA expression

The expressions of interleukin-1 (IL-1), tumor necrosis factor-α (TNF-α), cyclooxygenase-2 (COX-2) and Nuclear factor kappa-light-chain-enhancer of activated B cells (NF-κB) mRNA were significantly higher in the OC43 group than in the control group (Table 3, Figure 6). The expressions of IL-1 mRNA (sOC43: 27.75 ± 1.868; HGYGT: 21.92 ± 0.6333; p<0.05), TNF-α mRNA (OC43: 225.8 ± 24.08, HGYGT: 166.7 ± 8.045; p < 0.05), COX-2 mRNA (OC43: 10.89 ± 0.8811; HGYGT: 8.609 ± 0.3086; p < 0.05), and NF-κB mRNA (OC43: 6.482 ± 1.039, HGYGT: 3.807 ± 0.07861; p < 0.05) were significantly higher in cell infected with OC43 than in cells that were infected with OC43 and treated HGYGT (Table 3, Figure 6).

**Fig 6.**
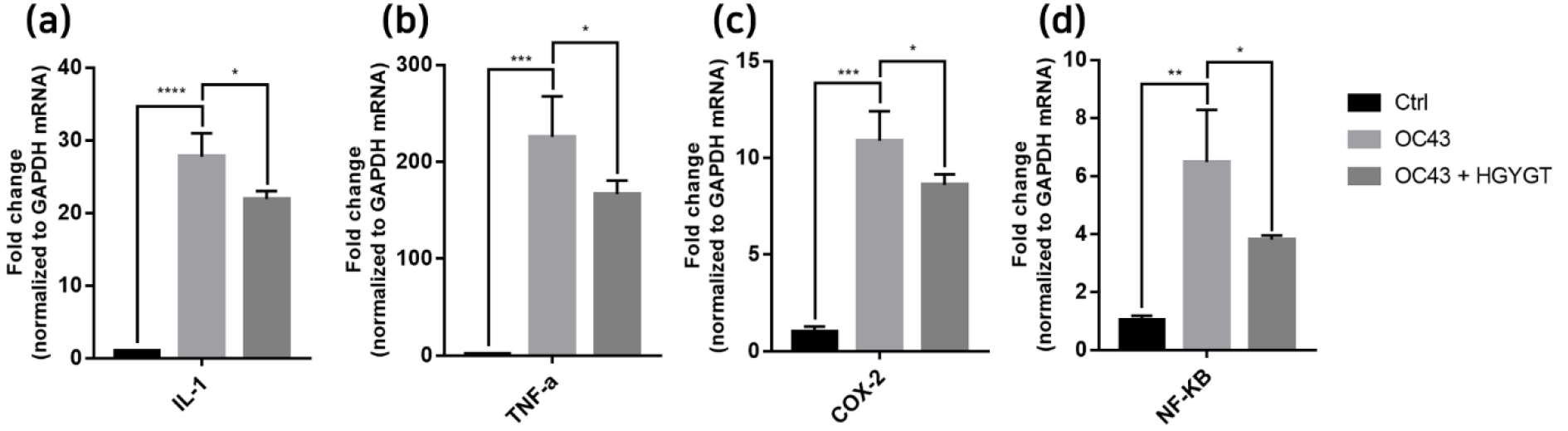
Pro-inflammatory cytokine mRNA expression in A549 cells. mRNA expression is measured using quantitative real-time PCR presented as mean ± S.D. (n= 3). Ctrl : cells with no treatment; OC43: cells treated with OC43; OC43 + HGYGT: cells treated with OC43 and HGYGT (100 μg/mL) (*; p<0.05, **<0.01, ***; p<0.001, ****; p<0.0001)

The expressions of Interferon Alpha Inducible Protein 6 (IFI6), IFI44, IFI44L, IFI27, Interferon Regulatory Factor 7 (IRF7), 2’-5’-Oligoadenylate Synthetase Like (OASL) and ISG15 mRNA were significantly higher in the OC43 group than in the control group (Table 3, Figure 7). The expressions of IFI6 mRNA (OC43: 150.5 ± 15.18; HGYGT: 84.61 ± 7.541; p < 0.01), IFI44 mRNA (OC43: 1413 ± 169.2; HGYGT: 761.7 ± 49.3; p < 0.05), IFI44L mRNA (OC43: 137.1 ± 4.453; HGYGT: 98.37 ± 2.426; p < 0.001), IFI27 mRNA (OC43: 506.2 ± 48.8; HGYGT: 328 ± 29.63; p <0.05), IRF7 mRNA (OC43: 7.599 ± 0.6307; HGYGT: 5.222 ± 0.7767; p < 0.05), OASL mRNA (OC43: 86.96 ± 1.575; HGYGT: 59.74 ± 2.015; p < 0.001) and ISG15 mRNA (OC43: 71.27 ± 4.874; HGYGT: 48.62 ± 4.098; p < 0.05) were significantly higher in cells that were only infected with OC43 than in infected cells treated HGYGT (Table 3, Figure 7).

**Fig 7.**
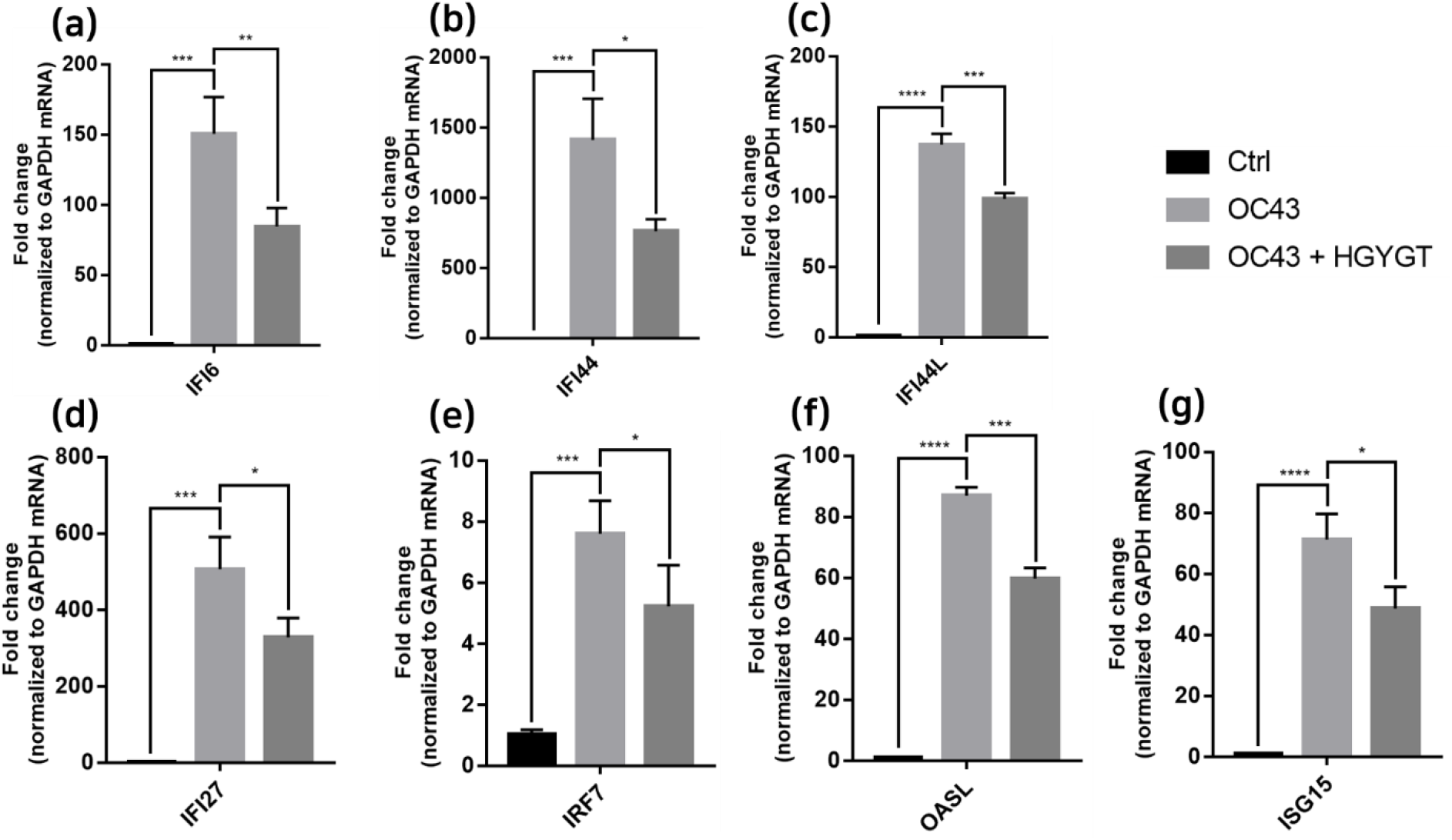
ISG mRNA expression in A549 cells. mRNA expression is measured using quantitative real-time PCR and presented as mean ± S.D. (n= 3). Ctrl : cells with no treatment; OC43: cells treated with OC43; OC43 + HGYGT: cells treated with OC43 and HGYGT (100 μg/mL) (*; p<0.05, **<0.01, ***; p<0.001, ****; p<0.0001)

The expressions of IκB kinase (IKK) and inducible nitric oxide synthase (iNOS) mRNA were significantly higher in the OC43 infected cells than in the control group (Table 3, Figure 8). The expressions of IKK mRNA (OC43: 2.6 ± 0.0503 HGYGT: 1.566 ± 0.1109; p<0.0001) and iNOS mRNA (OC43: 2.195 ± 0.05391; HGYGT: 1.372 ± 0.02985; p <0.0001) were significantly higher in the OC43 infected cells than in HGYGT treated infected cells (Table 3, Figure 8).

**Fig 8.**
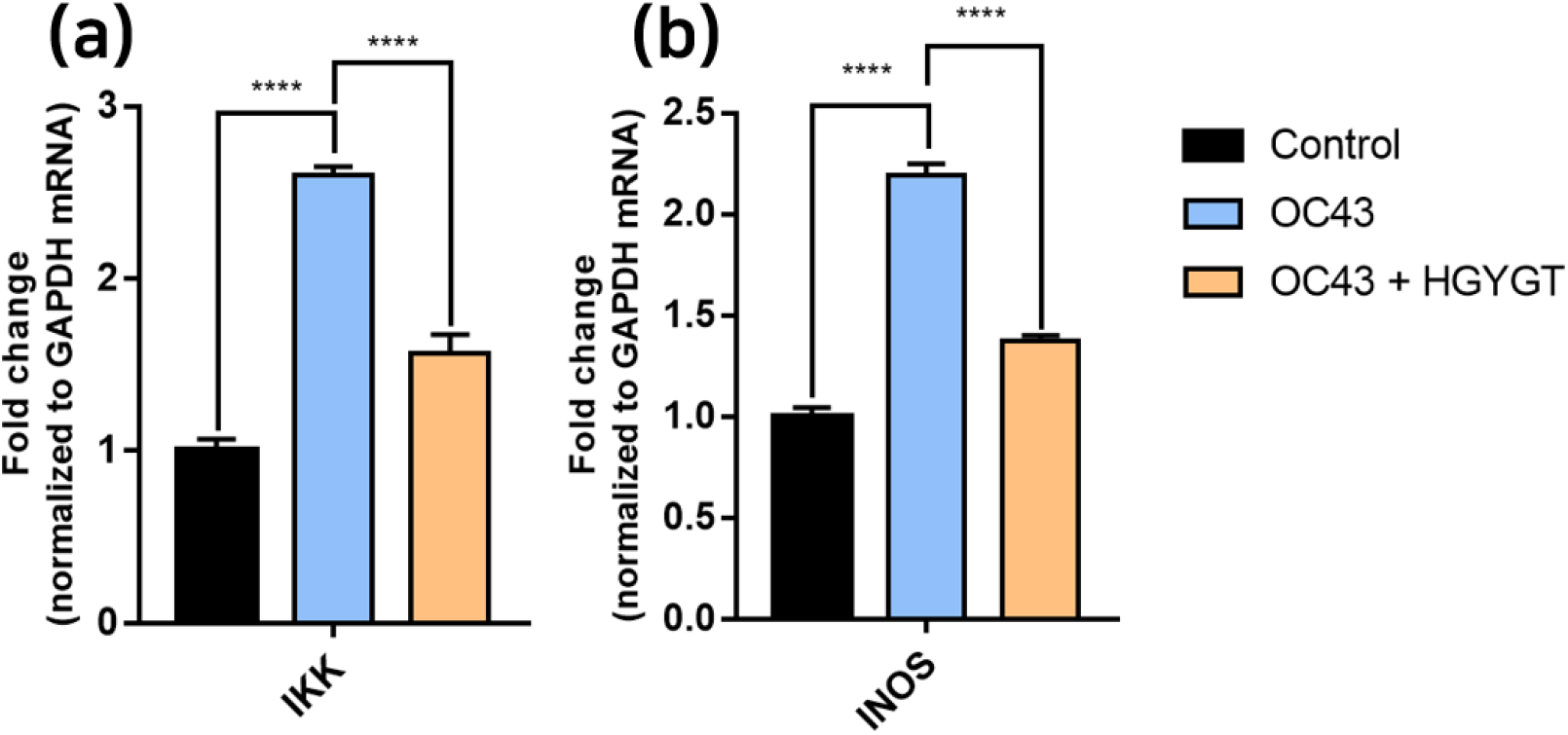
IKK, INOS mRNA expression in A549 cells. mRNA expression is measured using quantitative real-time PCR and presented as mean ± S.D. (n= 3). Ctrl : cells with no treatment; OC43: cells treated with OC43; OC43 + HGYGT: cells treated with OC43 and HGYGT (100 μg/mL) (****; p<0.0001)

## 4. Discussion

In December 2019, a number of pneumonia cases of unknown origin were diagnosed in Wuhan City, Hubei Province, China. From this pneumonia, a coronavirus with new genetic information, which has not yet been identified, was detected [5]. The World Health Organization named the virus SARS-CoV-2 as COVID-19 and declared a global pandemic [6]. From the beginning of the outbreak to March 23, 2021, 123,676,223 patients were infected with the virus worldwide, and 2,722,912 patients died.

SARS-CoV-2 belongs to the Coronaviridae family, the *Betacoronavirus* genus, and the *Sarbecovirus* subgenus. Betacoronaviruses include HCoV-OC43, HCoV-HKU1, SARS-CoV-1, MERS-CoV, and SARS-CoV-2. The SARS-CoV-1 virus infected more than 8,000 people and killed 774 in Guangdong, China from 2002 to 2003. The MERS-CoV virus infected 2,499 people and killed 861 in the Middle East from 2012 to 2019 [7,8].

SARS-CoV-2 is a positive-sense single-stranded RNA (ssRNA) virus, enclosed in an 80-220 nm envelope [9]. The virus recognizes a specific protein in the host cell membrane and binds to it, resulting in absorption by the cell. After the virus and the host cell are fused, the uncoated virus RNA invades the host cell. The RNA is replicated using host cell ribosomes and enzymes, and caspids are synthesized. Viruses formed by viral proteases are fused with the host cell membrane and released externally via budding (Figure 9). Antiviral drugs aim to phase out the life cycle of the virus to weaken or extinguish the invading virus by interfering with RNA transcription or inhibiting virus budding [10].

**Fig 9.**
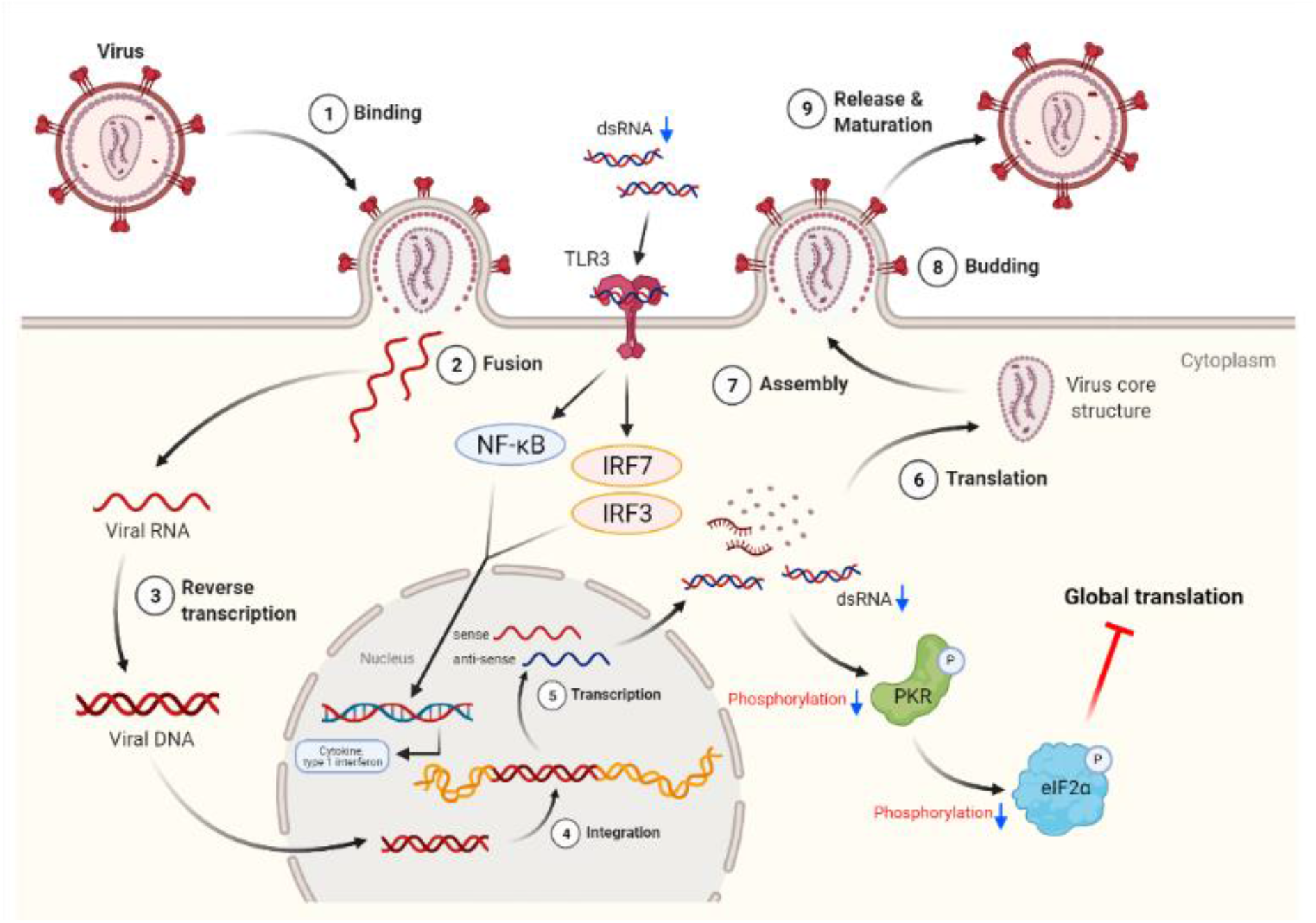
Antiviral effect of HGYGT on Coronavirus. (Created with BioRender.com)

Clinical signs of COVID-19 typically begin within one week after infection and include persistent fever, cough, nasal congestion, fatigue, and other symptoms of upper respiratory infections. Symptoms of the gastrointestinal tract have also been reported, and some cases are asymptomatic [11]. Symptoms may progress to difficulty breathing or extreme chest pain related to pneumonia, decreased oxygen saturation, and physiological changes observed through imaging such as chest X-ray or CT [12].

According to a study by Son et al.[13], 58.19% of Korean medicine doctors use uninsured herbal extracts. Insured herbal extracts are widely prescribed due to the low cost to patients, but differ from packaged herbal medications because they contain extracts of a single herb [14]. For this reason, Uninsured Herbal Extracts, which are most similar to packed herbal medicines, are widely prescribed despite the cost burden of patients. In this study, Uninsured Herbal Extracts were used for standardization and quality control of drugs.

HGYGT is listed in the Man-Byeong-Hoi-Chun of the Ming Dynasty, composed of 13 substances (Table 1). The HGYGT used in this study contains 0.5 g of *Schizonepeta tenuifolia Briquet, Scutellaria baicalensis Georgi, Saposhnikovia divaricata Schischkin, Angelica gigas Nakai, Poncirus trifoliata, Rafinesque, Paeonia lactiflora Pallas, Glycyrrhiza uralensis Fischer, Cnidium officinale Makino, Gardenia jasminoides Ellis, Forsythia suspensa Vahl* and 0.83 g of *Angelica dahurica Bentham et Hooker f, Platycodon grandiflorum A. De Candolle, Bupleurum falcatum Linné*. HGYGT is widely used to treat otolaryngeal diseases, including otitis media [15]. Previous studies regarding HGYGT have reported a genotoxicity evaluation [16], anti-allergic effects [17], anti-inflammatory effects through NF-κB activation control [18], antioxidant and anti-inflammatory effects through the reduction of iNOS and cytokine production [19], inhibition of inflammatory cell activation, and improvement of immune control [20].

PKR, also known as eukaryotic translation initiation factor 2-alpha kinase 2 (EIF2AK2), is autophosphorylated by dsRNA in infected cells, leading to an immune response [21]. pPKR is activated to inhibit global translation by preventing the transition of eIF2 to eIF2B [22]. Existing studies have reported that PKR is a signaling sensor for cellular metabolism [2]. The change in pPKR is proportional to the change in dsRNA. In this study, the expression of pPKR decreased as the amount of dsRNA decreased (Figure 2). Decrease of dsRNA expression inhibits the phosphorylation of PKR, thus inhibiting the activation of eIF2α. This reduces global translation and reduces protein synthesis (Figure 9). Recently, it has been reported that SARS-CoV-2 activates pPKR [23]. In this study, we found HGYGT could block phosphorylation of PKR with a decrease of virus RNA expression. This result signifies that HGYGT could be used as a drug to help the treatment for COVID-19.

We confirmed that IFI6, IFI44, IFI44L, IFI27, IRF7, OASL and ISG15 mRNA expressions were decreased in cells treated with HGYGT compared to those that were not treated with HGYGT (Figure 7). The reduced expressions of these proteins may affect the inactivation of type I Interferon (IFN) pathway. (Figure 9) Type 1 IFN pathway is a defense mechanism that induces an innate immune response, and excessive production of IFN-α/β can lead to the development of autoimmune diseases [24]. The reduction of ISGs that involved in type 1 IFN with HGYGT means it could be used for the treatment of various immune and inflammatory diseases. Previous studies on hepatitis C virus and RNA viruses have reported that IFI44, IFI44L and IRF7 are involved in translation [25]. In particular, IFI44L has been reported to be directly involved in the IFN response and acts as a regulator of antiviral responses [26]. It has been reported that IFI27 is involved in replication [27] and that OASL is involved in translation [25] and replication [28].

SARS-CoV-2 promotes excessive ROS. As ROS increases, H_2_O_2_ accumulates in tissues. The increase in ROS level promotes viral replication, causes oxidative inflammation, and cell apoptosis due to DNA damage. Recent studies have reported that catalase regulates cytokine in COVID-19, protects oxidative injuries, and inhibits replication of SARS-CoV-2 [29]. Viral infection increases free radicals, including nitrite oxide (NO), and depletes antioxidants [30]. RNA viruses cause oxidative stress, presumably due to the production of ROS in mitochondrial dysfunction, which occurs during virus infection with cells [31]. The production of iNOS involved in the oxidative injury involves activation of NF-kB/Rel [32]. Other studies have reported that inhibiting NF-kB pathways reduces iNOS mRNA and NO synthesis [33]. Both LPS and dsRNA are involved in type 1 IFN production. LPS and dsRNA activates antigen-presenting cells (APCs) via the Toll-like receptor (TLR) 3 and TLR4 signaling pathway [34]. Previous studies have reported the effect of HGYGT suppressing production of iNOS by reducing NF-kB/Rel in an LPS-induced oxidative injury [35]. Inactivated NF-kB/Rel fails to induce transcription at the NF-kB/Rel bond site in the TATA box of iNOS gene [36]. The expression of iNOS, IKK, NF-κB was decreased in cells treated with HGYGT compared to those that were not treated with HGYGT (Figure 8). Inflammation with OC43 increase IKK and iNOS mRNA levels in OC43 group. HGYGT reduces IKK activity, and it appears that NF-kB/Rel is inactivated due to reduced IκB phosphorylation, resulting in iNOS expression reduction. The reduction of IKK and iNOS mRNA levels after HGYGT treatment is believed to be likely to reduce levels of dsRNA and replication of coronavirus by acting on NF-kB/Rel pathways to protect oxidative injury.

The results of this study suggest that HGYGT may have antiviral effects. We evaluated the expression of several proteins and mRNAs to determine the mechanism of HGYGT. HGYGT reduces the expression of mRNAs for cytokines, such as IL-1, TNF-α, COX-2, NF-κB, iNOS, IKK and the expressions of mRNAs for ISGs involved in the type I IFN pathway, resulting in a decreased immune response and fewer clinical symptoms. We suggest that HGYGT of uninsured herbal extracts manufactured by pharmaceutical companies effectively reduces the virus via regulation in NF-kB/Rel pathways and reduces the symptoms of the coronavirus infection. However, our study is limited due to its in-vitro design and an insufficient sample size. More studies, including in-vivo and clinical studies, are needed to determine the effects of HGYGT on coronavirus infections.

## 5. Conclusions

HGYGT reduced the mRNA expression levels of coronavirus and inhibited the phosphorylation of PKR. It reduced mRNA expression level of iNOS and IKK. That means HGYGT may regulate NF-kB/Rel pathways and has antioxidative effect. HGYGT blocks a series of processes of ISG expression and reduces the inflammatory response in Beta coronavirus infected A549 cells. Thus, HGYGT has potential as a therapeutic agent for various clinical symptoms caused by coronavirus.

## Abbreviations

APC: antigen-presenting cells
COVID-19: coronavirus disease 2019
COX-2: Cyclooxygenase-2
dsRNA: Double-stranded RNA
eIF2: Eukaryotic initiation factor 2
EIF2AK2: Eukaryotic translation initiation factor 2-alpha kinase 2
HCoV: Human CoV
HGYGT: Hyeonggae Yeongyo-tang
IFI: Interferon Alpha Inducible Protein
IFN: Interferon
IKK: IκB kinase
IL: Interleukin
iNOS: inducible nitric oxide synthase
IRF: Interferon Regulatory Factor
ISGs: Interferon stimulated genes
MERS: Middle East respiratory syndrome
NF-κB: Nuclear factor kappa-light-chain-enhancer of activated B cells
NO: nitrite oxide
OASL: 2’-5’-Oligoadenylate Synthetase Like
PKR: Protein kinase RNA-activated
ROS: Reactive Oxygen Species
RT: Reverse transcription
SARS-CoV: severe acute respiratory syndrome coronavirus
ssRNA: single-stranded RNA
TLR: Toll-like receptor
TNF-α: Tumor necrosis factor-α

## Data Availability

The dataset generated or analyzed in this study are available from the corresponding author upon reasonable request.

## Conflicts of Interest

The authors declare that there is no conflict of interest regarding the publication of this article

## Acknowledgments

We are grateful to the experts for their valuable advice.

## Notes

### Competing Interest Statement

The authors have declared no competing interest.

## References

[1] O.J. McElvaney, N.L. McEvoy, O.F. McElvaney et al., “Characterization of the Inflammatory Response to Severe COVID-19 Illness,” Am J Respir Crit Car Med, vol. 202, no.6, pp. 812–821, 2020.

[2] M.A. García, E.F. Meurs and M. Esteban, “The dsRNA protein kinase PKR: Virus and cell control,” Biochimie, vol. 89, no.6-7, pp. 799–811, 2007.

[3] S.J. Poynter and S.J. DeWitte-Orr, “Understanding Viral dsRNA-Mediated Innate Immune Responses at the Cellular Level Using a Rainbow Trout Model,” Front Immunol, vol. 9, pp. 829, 2018.

[4] M.L Reshi, Y.C Su, and J.R Hong, “RNA Viruses: ROS-Mediated Cell Death,” International Journal of Cell Biology, vol. 2014, Article ID 467452, 2014.

[5] D. Paraskevis, E.G. Kostaki, G. Magiorkinis et al., “Full-genome evolutionary analysis of the novel corona virus (2019-nCoV) rejects the hypothesis of emergence as a result of a recent recombination event,” Infect Genet Evol, vol. 79, Article ID 104212, 2020.

[6] World Health Organization, “WHO Director-General’s remarks at the media briefing on 2019-nCoV on 11 February 2020 [Internet],” Geneva: World Health Organization; 2020.

[7] J. Guarner, “Three Emerging Coronaviruses in Two Decades,” Am J Clin Pathol, vol 153, no. 4, pp. 420–421, 2020.

[8] T.T Yao, J.D. Qian, W.Y. Zhu, Y. Wang and G.Q. Wang, “A systematic review of lopinavir therapy for SARS coronavirus and MERS coronavirus—A possible reference for coronavirus disease-19 treatment option,” J Med Virol, vol 92, no. 6, pp.556–563, 2020.

[9] J.A. Englund, Y.J. Kim and K. McIntosh, “Human coronaviruses, including Middle East respiratory syndrome coronavirus,” In: J. Cherry, G.J. Demmler Harrison, S.L. Kaplan, W.J. Steinbach and P.J. Hotez editors, “Feigin and Cherry’s textbook of pediatric infectious disease,” 8th ed. Philadelphia : Elsevier Inc, pp. 1846–1854, 2019.

[10] J.M. Parks and J.C. Smith, “How to Discover Antiviral Drugs Quickly,” N Engl J Med, vol. 382, pp. 2261–2264, 2020.

[11] W. Guan, Z. Ni, Y. Hu et al., “Clinical characteristics of 2019 novel coronavirus infection in China,” N Engl J Med, vol. 382, pp. 1708–1720, 2020.

[12] T.P Velavan and C.G. Meyer, “The COVID‐19 epidemic,” Trop Med Int Health, vol. 25, no. 3, pp. 278–280, 2020.

[13] C.H. Son, Y.H. Kim and S. Lim, “A Study on Korean Oriental Medical Doctors’ Use of Uninsured Herbal Extracts and How to Promote the Insurance Coverage of Such Herbal Extracts,” J Korean Oriental Med, vol. 30, no. 4, pp. 64–78, 2009.

[14] S.J. Park, K.S. Kim, H.S. Kim et al., “A quantitative analysis of marker compounds in single herb extracts by the standard of KHP,” Kor J Herbology, vol. 29, no. 3, pp. 35–42, 2014.

[15] M.R. Yang, K.S. Jin, H.J. Lee, M.W. Kwon and E.J. Park, “A Clinical study on the Therapeutic effect of Kamihyunggyeyungyotang for Pediatric Recurrent Otitis Media with Effusion,” J Korean Oriental Pediatrics, vol. 15, no. 2, pp. 87–100, 2001.

[16] S.Y. Jee, S.Y. Hwang, J.R. Lee and S.C. Kim, “A study of Genotoxicity Test of Hyeong-gae-yeon-gyo-tang extract,” Kor J Herbology, vol. 22, no.4, pp. 287–300, 2007.

[17] T.S Yoo, Y.S Chin and K.M Jeong, “Study of the effects of Hyunggaeyeungyotang on the Anti-allergic effect in rats and mice,” The Journal of Pediatrics of Korean Medicine, vol. 4, no. 1, pp.19–30, 1990.

[18] J.H. Park and S.U. Hong, “The Effects of Hyunggaeyungyo-tang of Suppression of iNOS Production on Mice with Allergic Rhinitis,” The Journal of Korean Oriental Medical Ophthalmology & Otolaryngology & Dermatology, vol. 25, no.1, pp.12–21, 2012.

[19] M.J. Kim, J.R. Lee, S.C. Kim and S.Y. Jee, “Inhibitory Effect of Hyeonggaeyeongyo-tang Water Extract on production of Nitric Oxide, IL-6 and Expression of iNOS, COX-2 in LPS - Activated Raw 264.7 Cells,” Korean Journal of Oriental Physiology & Pathology, vol. 21, no. 2, pp. 491–497, 2007.

[20] R.Y. Kang, B.K. Park, S.B. Kim, H.J. Choi and D.H. Kim, “The effects of HYT on various immunological factors related to pathogenesis of allergic dermatitis in NC/Nga mice induced by Biostir AD,” Journal of Haehwa Medicine, vol. 18, no. 2, pp. 47–62, 2009.

[21] R.C. Patel, P. Stanton, N.M. McMillan, B.R. Williams and G.C. Sen, “The interferon-inducible double-stranded RNA-activated protein kinase self-associates in vitro and in vivo,” Proc Natl Acad Sci USA, vol. 92, no. 18, pp. 8283–8287, 1995.

[22] E.F. Meurs, Y. Watanabe, S. Kadereit et al., “Constitutive expression of human double-stranded RNA-activated p68 kinase in murine cells mediates phosphorylation of eukaryotic initiation factor 2 and partial resistance to encephalomyocarditis virus growth,” J Virol, vol. 66, pp. 5805–5814, 1992.

[23] Y. Li, D.M. Renner, C.E. Comar et al., “SARS-CoV-2 induces double-stranded RNA-mediated innate immune responses in respiratory epithelial-derived cells and cardiomyocytes,” PNAS, vol. 118, no. 6, e2022643118, 2021.

[24] J. Banchereau and V. Pascual, “Type ? interferon in systemic lupus erythematosus and other autoimmune disease,” Immunity, vol. 25, no. 3, pp. 383–392, 2006.

[25] J.W. Schoggins, S.J. Wilson, M. Panis et al., “A diverse range of gene products are effectors of the type I interferon antiviral response,” Nature vol. 472, pp. 481–485, 2011.

[26] M.L. DeDiego, L. Martinez-Sobride and D.J. Topham, “Novel Functions of IFI44L as a Feedback Regulator of Host Antiviral Responses,” J Virol, vol. 93, no. 21, e01159–19, 2019.

[27] Y. Itsui, N. Sakamoto, M. Kurosaki et al., “Expressional screening of interferon stimulated genes for antiviral activity against hepatitis C virus replication,” J Viral Hepat, vol. 13, no. 10, pp. 690–700, 2006.

[28] M. Ishibashi, T. Wakita and M. Esumi, “2’,5’-Oligoadenylate synthetaselike gene highly induced by hepatitis C virus infection in human liver is inhibitory to viral replication in vitro,” Biochem Biophys Res Commun, vol. 392, no. 3, pp. 397–402, 2010.

[29] M. Qin, Z. Cao, J. Wen et al., “An Antioxidant Enzyme Therapeutic for COVID-19,” Adv. Mater, vol. 32, e2004901, 2020.

[30] F.C. Camini, C.C. da Silva Caetano, L.T. Almeida and C.L de Brito Magalhães, “Implications of Oxidative Stress on Viral Pathogenesis,” Arch Virol., vol. 162, no. 4, pp. 907–917, 2017.

[31] A.V. Ivanov, V.T. Valuev-Elliston, O.N. Ivanova et al., “Oxidative Stress during HIV Infection: Mechanisms and Consequences,” Oxid Med Cell Longev, vol. 2016, e8910396, 2016.

[32] Q.W. Xie, Y. Kashiwabara and C. Nathan, “Role of transcription factor NF-kappa B/Rel in induction of nitric oxide synthase,” Journal of Biological Chemistry, vol. 269, no. 7, pp. 4705–4708, 1994.

[33] M. Matsumura, H. Kakishita, M. Suzuki, N. Banba and Y. Hattori, “Dexamethasone suppresses iNOS gene expression by inhibiting NF-κB in vascular smooth muscle cells,” Life Sci., vol. 69, no.9, pp. 1067–1077, 2001.

[34] K. Hoebe and B. Beutler, “LPS, dsRNA and the interferon bridge to adaptive immune responses: Trif, Tram, and other TIR adaptor proteins,” J Endotoxin Res., vol. 10, no. 2, pp. 130–136, 2004.

[35] J.H. Park, J.C. Kim and S.U. Hong, “The effects of Hyunggaeyungyo-tang of suppression of iNOS production on RAW 264.7 cell,” The Journal of Korean Oriental Medical Ophthalmology & Otolaryngology & Dermatology, vol. 24, no. 1, pp. 78–85, 2011.

[36] C.J. Lowenstein, E.W. Alley, P. Raval et al., “Macrophage nitric oxide synthase gene: two upstream regions mediate induction by interferon gamma and lipopolysaccharide,” Proc Natl Acad Sci USA, vol. 90, no. 20, pp. 9730–9734, 1993.

